# ATG deficiency impairs stationary-phase microlipophagy through acetic acid-induced clustering of Niemann–Pick type C proteins

**DOI:** 10.64898/2026.04.22.720228

**Authors:** Takuma Tsuji, Megumi Fujimoto, Nobuo N. Noda, Toyoshi Fujimoto

## Abstract

While the role of autophagy-related (ATG) proteins in microautophagy remains unclear, their absence in budding yeast has been reported to impair stationary-phase microlipophagy. Here, we show that this defect in ATG-deficient (*atg*Δ) cells arises not from a direct requirement of ATG proteins for the execution of microlipophagy but from accumulation of acetic acid (AA) in the medium. High concentrations of AA in the medium of *atg*Δ cells trigger the clustering of Niemann–Pick type C (NPC) proteins, causing impairment of raft-like vacuolar microdomain formation and suppression of microlipophagy. Lowering extracellular AA rapidly dissolves NPC protein clusters, restores vacuolar microdomains, and rescues microlipophagy in *atg*Δ cells. Conversely, elevating AA concentrations in the medium of wild-type cells induces NPC protein clusters and microlipophagy defects. These findings demonstrate that stationary-phase microlipophagy can proceed independently of ATG proteins and that the defect in *atg*Δ cells can be rescued by normalizing extracellular AA levels.

## Introduction

Autophagy plays a crucial role in recycling cellular components and maintaining homeostasis.^1^ During macroautophagy, substrates are enwrapped by isolation membranes to form autophagosomes, which then fuse with the lysosome for degradation.^2^ By contrast, during microautophagy, substrates are directly engulfed by inward invagination of the lysosomal/vacuolar membrane.^3,4^ Microautophagy is conserved across diverse species, from yeasts and plants^5^ to mammals.^6–8^ It facilitates the degradation of various cellular components, including the nucleus,^9^ endoplasmic reticulum (ER),^10–12^ lipid droplets (LDs),^13–15^ mitochondria,^16^ peroxisomes,^17,18^ and vacuolar membrane proteins.^19^ Despite recent advances in elucidating the molecular mechanisms of autophagy, the precise role of ATG proteins in microautophagy remains poorly defined.^3,4,20,21^

This gap in knowledge is particularly evident for lipid droplet (LD) microautophagy (microlipophagy) in the budding yeast *Saccharomyces cerevisiae*. Microautophagy in budding yeast has been observed under various conditions, including nitrogen starvation,^14^ glucose starvation,^22^ stationary phase,^15^ post-diauxic shift phase,^23^ ethanol-depleted phase,^24^ and ER stress.^25,26^ Although ATG proteins have been reported to be necessary for microlipophagy induced by nutrient starvation or during the stationary phase,^14,15,22^ they have been found dispensable for microlipophagy during the post-diauxic shift, ethanol depletion, or ER stress.^23–26^ Given that inward invagination of the vacuolar membrane is a common feature of microlipophagy across these conditions, such differences raise questions as to whether and how ATG proteins are involved in microlipophagy.

Microlipophagy in stationary-phase yeast proceeds through the formation of raft-like vacuolar microdomains.^27,28^ We previously showed that two Niemann–Pick type C (NPC) proteins, Ncr1 and Npc2, which collaborate to transport free sterol to the vacuolar membrane, are essential for microdomain formation.^28^ Ncr1 and Npc2 are normally distributed in the membrane and in the lumen of the vacuole,^29–31^ respectively, but in ATG-deficient (*atg*Δ) cells in stationary phase, they co-accumulate in a small number of puncta (hereafter referred to as NPC puncta).^28^ We hypothesized that this abnormal distribution of NPC proteins underlies the microlipophagy defects in *atg*Δ cells and that restoring their proper localization could rescue microlipophagy.

In this study, we tested this hypothesis by analyzing the factors responsible for NPC puncta formation and found that these puncta arise in response to elevated acetic acid (AA) concentrations in the medium. The NPC puncta behaved as reversible clusters with liquid-like properties and rapidly dissolved when the AA concentration was reduced by medium exchange. Lowering extracellular AA not only normalized NPC protein distribution but also restored vacuolar microdomain formation and microlipophagy in *atg*Δ cells. Conversely, elevating AA levels in wild-type (WT) cell cultures induced NPC puncta and impaired microlipophagy. These results indicate that ATG proteins are not directly required for the execution of stationary-phase microlipophagy but rather act indirectly by maintaining proper metabolism to prevent AA accumulation.

## Results

### Formation of NPC puncta is induced by the medium of stationary-phase *atg*Δ cells

In WT cells, Ncr1-yeGFP and Npc2-yeGFP were distributed in the vacuolar membrane and in the vacuolar lumen, respectively, both in the logarithmic growth phase and stationary phase (Figs. 1A and S1A). By contrast, in *atg7*Δ cells, Ncr1 and Npc2 co-clustered to form puncta in the stationary phase, whereas their distribution in the logarithmic growth phase was similar to that in WT cells (Figs. 1A and S1A).^28^ We serendipitously found that these NPC puncta in stationary-phase *atg7*Δ cells dissolved within several minutes after transfer to fresh medium (Fig. S1B). NPC puncta also dissolved when cells were transferred to the stationary-phase WT medium or to distilled water (DW) (Figs. 1B, S1C, and S1D). NPC puncta observed in stationary-phase *atg1*Δ, *atg3*Δ, *atg5*Δ, and *atg18*Δ cells also disappeared by the same treatment (Fig. S1E). These results suggest that some factors present in the stationary-phase *atg*Δ medium, but not in WT medium, may drive NPC puncta formation. Supporting this idea, when *atg7*Δ cells that had been rinsed with DW were returned to their original stationary-phase medium, the NPC puncta reappeared (Figs. 1C and S1F). Furthermore, exposure of WT cells to the stationary-phase *atg7*Δ medium induced NPC puncta formation, and subsequent washing with DW reversed this effect (Figs. 1D and 1E). These findings show that the distribution of Ncr1 and Npc2 can be manipulated by changing the culture medium.

**Figure 1.**
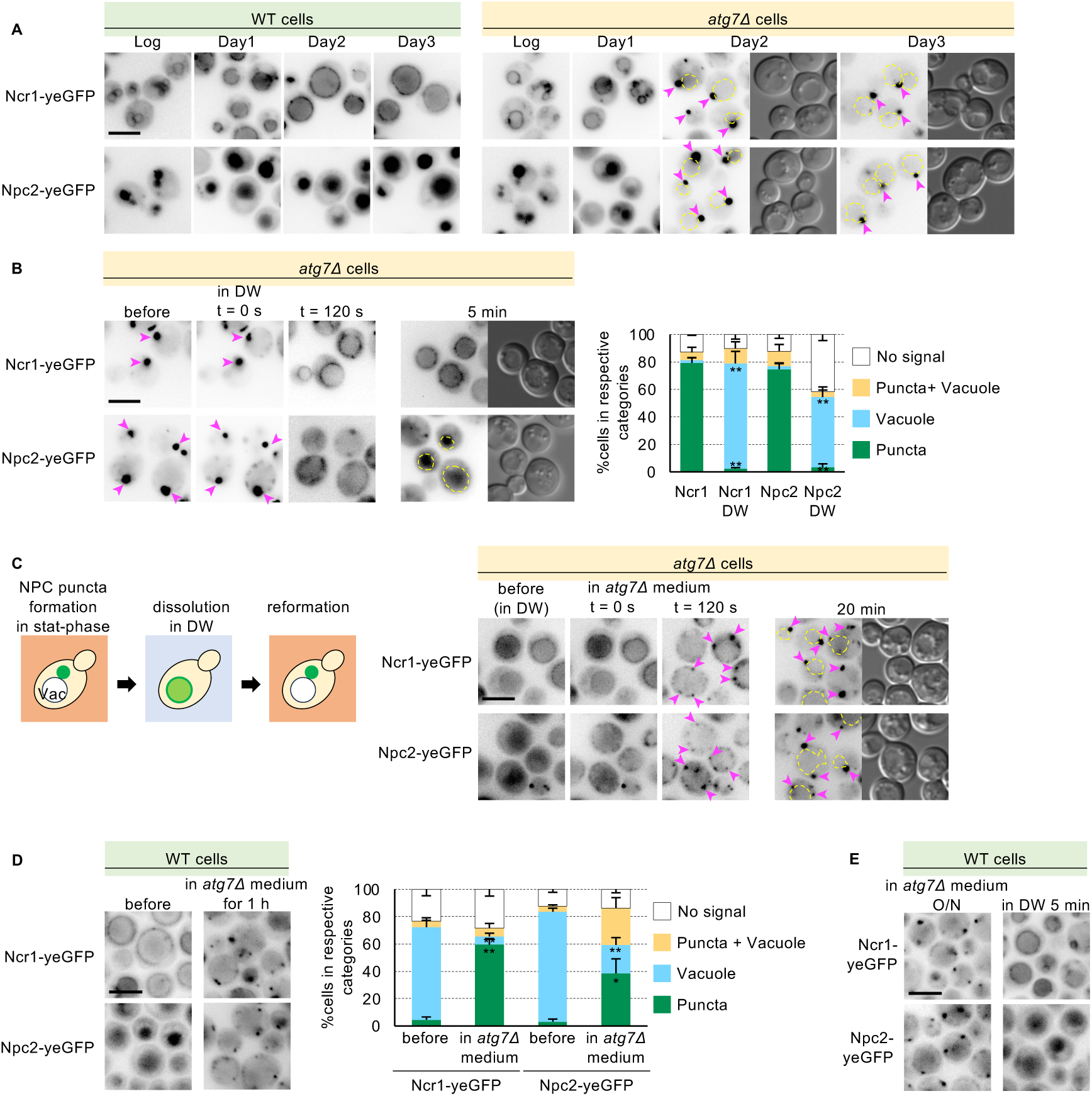
Formation of NPC puncta is induced by the medium of stationary-phase *atg*Δ cells. A) Ncr1-yeGFP and Npc2-yeGFP localization in log- and stationary-phase WT and *atg7*Δ cells. Cells were diluted in fresh medium to OD_600_ = 0.15 on Day 0 and imaged during log phase, on Day 1, and during stationary phase (Days 2 and 3). Arrowheads indicate NPC puncta. Bar, 5 μm. B) Time-lapse images of Ncr1-yeGFP and Npc2-yeGFP in stationary-phase *atg7*Δ cells after DW washing. Arrowheads indicate NPC puncta. Bar, 5 μm. The graph shows the proportions of cells exhibiting punctate, vacuolar, or mixed distributions 5 min after washing with DW. For each condition, more than 50 cells were counted per experiment, and the mean ± SEM of three independent experiments is shown. Statistical significance between DW-washed and non-washed cells was assessed by Student’s t-test; \*\**p* < 0.01. C) Time-lapse images of Ncr1-yeGFP and Npc2-yeGFP in stationary-phase *atg7*Δ cells that were washed with DW and returned to the original medium. Arrowheads indicate NPC puncta. Bar, 5 μm. D) NPC puncta formed in stationary-phase WT cells that were cultured for 1 h in the stationary-phase *atg7*Δ medium. Bar, 5 μm. The graph shows the proportions of cells showing punctate, vacuolar, or mixed distributions after more than 1 h in the stationary-phase *atg7*Δ medium. Data represent mean ± SEM (*n* = 3, >50 cells/condition; \*\**p* < 0.01, \**p* < 0.05 vs. before-treatment cells, Student’s t-test). E) Stationary-phase WT cells were cultured overnight in the stationary-phase *atg7*Δ medium and then washed with DW. Bar, 5 μm.

### Acetic acid drives reversible NPC puncta formation

As a candidate factor enriched in the stationary-phase *atg*Δ medium that induces NPC puncta, we focused on AA, which has been reported to accumulate during stationary phase and to increase abnormally in *atg*Δ medium.^32,33^ Consistent with these reports, the stationary-phase media of *atg1*Δ and *atg7*Δ cells contained more than twice as much AA as WT medium (Fig. 2A).

**Figure 2.**
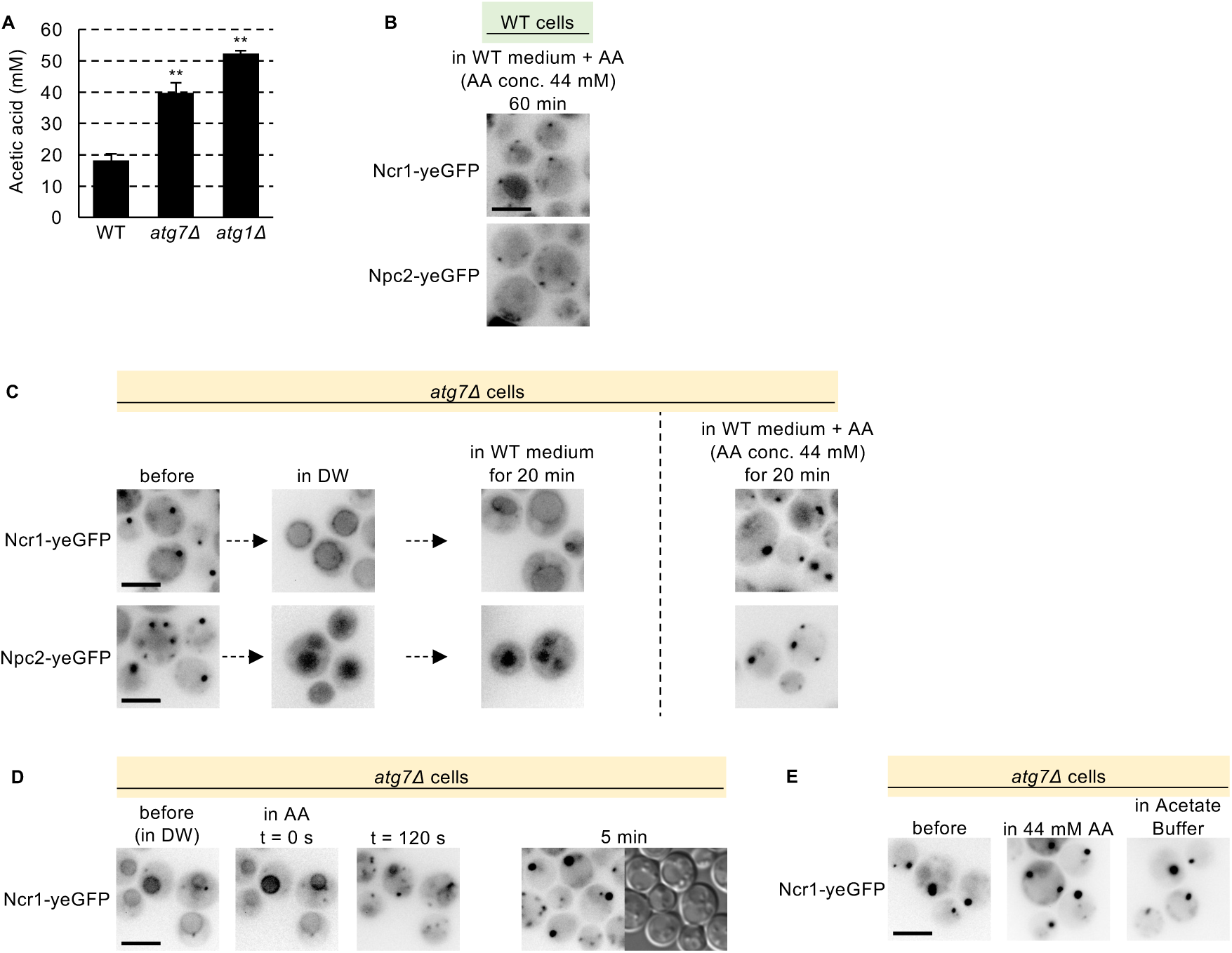
Acetic acid drives reversible NPC puncta formation. A) AA concentrations in the medium of stationary-phase WT, *atg7*Δ, and *atg1*Δ cells. Data represent mean ± SEM (*n* = 3; \*\**p* < 0.01 vs. WT cells, Student’s t-test). B) Micrographs of stationary-phase WT cells expressing Ncr1-yeGFP or Npc2-yeGFP cultured in WT medium supplemented with AA. Bar, 5 μm. C) Micrographs of stationary-phase *atg7*Δ cells expressing Ncr1-yeGFP or Npc2-yeGFP washed with DW and subsequently cultured in WT medium with or without AA supplementation. Bar, 5 μm. D) 44 mM AA solution induces NPC puncta in DW-washed *atg7*Δ cells. Time-lapse images and representative images taken after 5 min, showing reformation of NPC puncta. Bar, 5 μm. E) Washing *atg7*Δ cells with 44 mM AA solution or 50 mM acetate buffer (pH 3.6) maintains NPC puncta. Bar, 5 μm.

We therefore tested whether elevated AA is sufficient to induce NPC puncta. Increasing the AA concentration of WT medium to 44 mM, a concentration measured in the stationary-phase *atg7*Δ medium, induced NPC puncta in WT cells (Fig. 2B). Similarly, when NPC puncta in *atg7*Δ cells were first dissolved by DW washing, they reformed upon transfer to WT medium containing 44 mM AA, but not upon transfer to WT medium without AA supplementation (Fig. 2C). Importantly, NPC puncta also reformed in DW-washed *atg7*Δ cells upon incubation in DW supplemented with 44 mM AA (Fig. 2D). These results indicate that elevated AA is sufficient to induce NPC puncta.

Because the pKa of AA is 4.76, most AA in the stationary-phase medium (pH ∼3) is undissociated (Fig. S2A). We therefore tested whether extracellular undissociated AA is required to maintain NPC puncta. In contrast to the rapid dissolution observed upon DW washing (Fig. 1B), NPC puncta in *atg7*Δ cells did not dissolve upon washing with either 44 mM AA or 50 mM acetate buffer (pH 3.6) (Fig. 2E), which were calculated to contain 43 mM and 47 mM undissociated AA, respectively, according to the Henderson–Hasselbalch equation. Consistently, adding NaOH to raise the medium pH above 4.76 eliminated NPC puncta in *atg7*Δ cells, whereas adding NaCl at the same molar concentration had no effect (Fig. S2B). Furthermore, NPC puncta dissolved immediately when cells were washed with DW acidified to pH 3.0 with HCl, indicating that protons alone are insufficient to maintain NPC puncta (Fig. S2C). This effect was also not a general property of organic acids, because fumaric, succinic, citric, and malic acids failed to maintain NPC puncta under the conditions tested (Fig. S2D). Together, these results demonstrate that high extracellular levels of undissociated AA are required to maintain NPC puncta.

Because AA exposure is known to cause cytosolic acidification,^34^ we tested whether NPC puncta formation is driven by a drop in bulk cytosolic pH. However, treatment of WT cells with the stationary-phase *atg7*Δ medium induced NPC puncta (Fig. 1D) without altering bulk cytosolic pH (Fig. S2E). Additionally, although undissociated AA can enter cells either by free diffusion or via Fps1,^35,36^ NPC puncta still formed in *atg7*Δ*fps1*Δ cells, indicating that Fps1-mediated transport is not required (Fig. S2F). Together, these findings indicate that AA-induced NPC puncta formation does not require Fps1-mediated uptake or a measurable decrease in bulk cytosolic pH.

### Acetic acid suppresses microlipophagy by inhibiting vacuolar microdomain formation

The above results led us to hypothesize that elevated AA levels, rather than the absence of ATG proteins per se, underlie impaired microlipophagy in stationary-phase *atg*Δ cells. To test this hypothesis, we first asked whether reducing AA restores vacuolar microdomain formation in *atg7*Δ cells, using Vph1-mRFP as a marker,^27^ because vacuolar microdomains are essential for LD uptake during microlipophagy.^28^ Prior to the DW wash, Ncr1-yeGFP was distributed in puncta, while Vph1-mRFP was uniformly distributed throughout the vacuolar membrane (Fig. 3A). Upon DW washing, Ncr1-yeGFP redistributed from puncta to the vacuolar membrane, and vacuolar microdomains excluding both Vph1-mRFP and Ncr1-yeGFP formed (Fig. 3A). Similarly, reducing AA by transferring *atg7*Δ cells to WT medium, which contains less AA than stationary-phase *atg7*Δ medium, restored microdomain formation, whereas washing the cells with AA did not (Fig. 3B).

**Figure 3.**
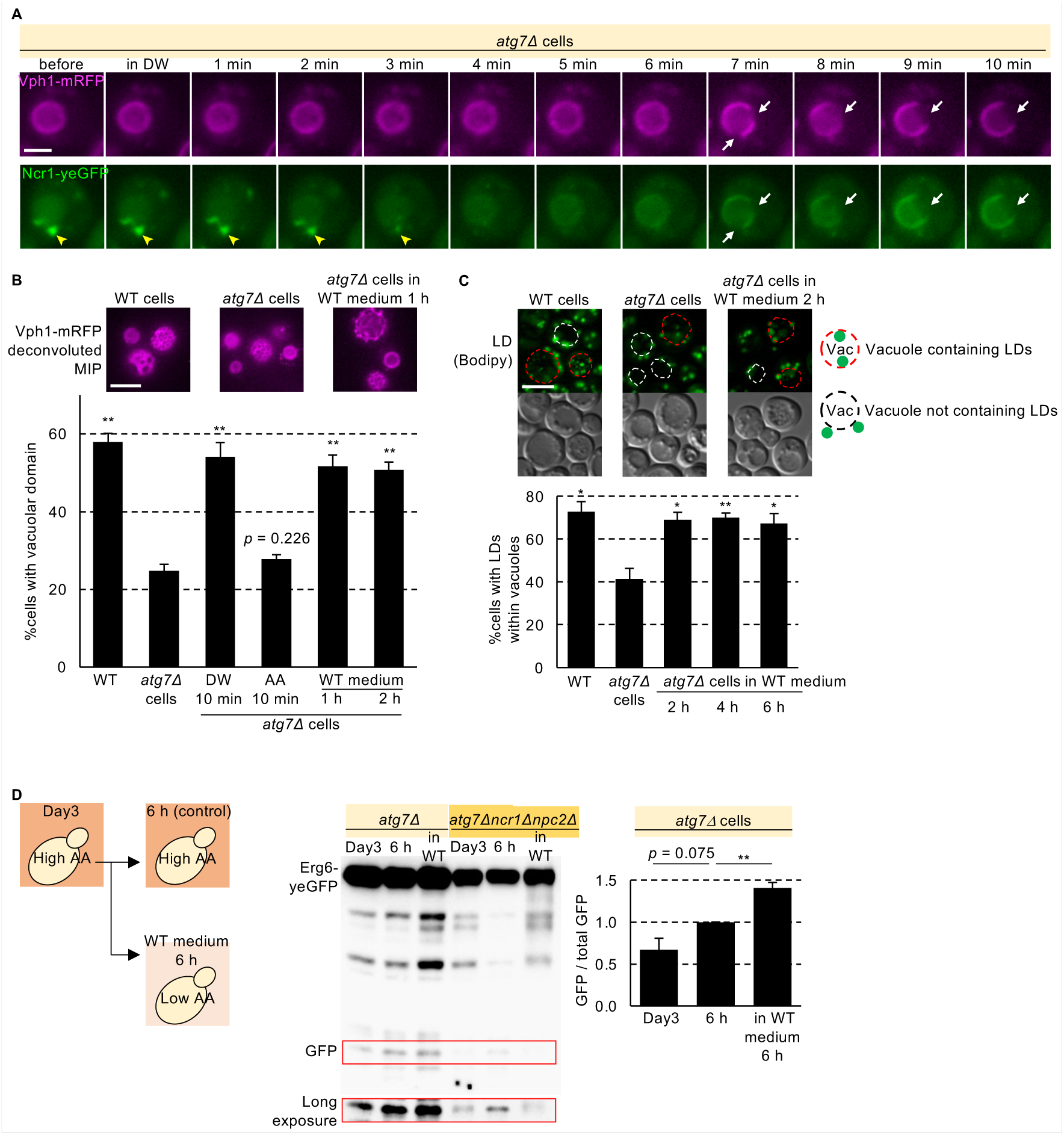
Acetic acid suppresses vacuolar microdomain formation and microlipophagy. A) Time-lapse images of stationary-phase *atg7*Δ cells expressing Ncr1-yeGFP and Vph1-mRFP after DW washing. At 4 min after washing, the NPC puncta (yellow arrowhead) dissolved. At 7 min, vacuolar membrane microdomains formed (white arrows). Bar, 2.5 μm. B) Deconvoluted maximum-intensity projection images of Vph1-mRFP in WT and *atg7*Δ cells in the stationary phase, as well as *atg7*Δ cells cultured in the stationary-phase WT medium for 1 h. Nine optical sections, each 300 nm thick, were used. Bar, 5 μm. The graph represents the proportions of cells with vacuolar microdomains. Data represent mean ± SEM (*n* = 3, >50 cells/condition; \*\**p* < 0.01 vs. *atg7*Δ cells, Student’s t-test). C) Micrographs of BODIPY-stained LDs in WT, *atg7*Δ, and *atg7*Δ cells cultured in the stationary-phase WT medium. The red dotted lines indicate vacuoles containing LDs, whereas the white dotted lines indicate vacuoles without LDs. Bar, 5 μm. The graph shows the proportions of cells with LDs within the vacuoles. Data represent mean ± SEM (*n* = 3, >50 cells/condition; \*\**p* < 0.01, \**p* < 0.05 vs. *atg7*Δ cells, Student’s t-test) D) Vacuolar degradation of Erg6-yeGFP in *atg7*Δ and *atg7*Δ*ncr1*Δ*npc2*Δ cells cultured for 6 h in the stationary-phase WT medium. Data represent mean ± SEM (*n* = 3; \*\**p* < 0.01 vs. 6 h (control), Student’s t-test)

Because lowering AA restores vacuolar microdomain formation in *atg7*Δ cells, we next asked whether it also restores microlipophagy during stationary phase. To test this, *atg7*Δ cells on Day 3 of stationary phase were transferred to WT medium and analyzed 6 h later. Under these conditions, the proportion of cells with LDs in the vacuole increased, and vacuolar degradation of the LD marker Erg6-yeGFP increased (Figs. 3C, 3D, S3A, and S3B). This degradation required NPC proteins, consistent with their role in microdomain formation (Figs. 3D, S3A and S3B).^28^ Together, these results suggest that lowering AA restores microlipophagy in *atg7*Δ cells.

The increased proportion of *atg7*Δ cells with LDs in the vacuole after transfer to WT medium suggests de novo induction of microlipophagy. However, because a substantial fraction of *atg7*Δ cells already contained vacuolar LDs on Day 3 (Fig. 3C), the increase in Erg6-yeGFP degradation observed 6 h later could reflect, at least in part, continued degradation of LDs that had entered the vacuole before Day 3, rather than de novo induction of microlipophagy after AA reduction. To distinguish between these possibilities, we established a controlled AA-shift assay. AA was added on Day 1 (24 h after cultivation began), and on Day 3 the medium was replaced with stationary-phase WT medium without additional AA supplementation. In *atg7*Δ cells, live-cell imaging before lowering AA on Day 3 showed that vacuoles were largely devoid of LDs (Movie 1). When AA levels were lowered on Day 3, Erg6-yeGFP degradation increased by Day 4 compared with cells kept in high-AA medium (Fig. S3C). Live imaging also showed LD uptake into the vacuole after the AA reduction (Movies 1 and 2). These results indicate that AA reduction is sufficient to induce microlipophagy in *atg7*Δ cells, rather than simply enhancing ongoing microlipophagic degradation initiated before Day 3. We next asked whether AA levels also reversibly control stationary-phase microlipophagy in WT cells. As in *atg7*Δ cells, vacuoles in WT cells exposed to AA from Day 1 were largely devoid of LDs before the AA reduction on Day 3 (Movie 1). Reducing AA on Day 3 increased Erg6-yeGFP degradation by Day 4, and live-cell imaging also showed LD uptake into the vacuole (Fig. S3C and Movies 1and 3). These findings indicate that microlipophagy is reversibly controlled by AA levels in both WT and *atg7*Δ cells. Together, our results show that elevated AA levels impair vacuolar microdomain formation and thereby suppress microlipophagy, whereas lowering AA restores both processes even in the absence of ATG proteins.

### NPC proteins form reversible clusters with liquid-like properties in vacuolar protrusions

To characterize the morphology and biophysical properties of NPC puncta, we first investigated their subcellular location using quick-freezing and freeze-fracture replica labeling electron microscopy (QF-FRL). QF-FRL analysis with labeling for phosphatidylinositol 3-phosphate revealed that vacuolar membranes protrude into the cytoplasm in stationary-phase *atg7*Δ cells (Fig. 4A). Ncr1-yeGFP, expressed either from the endogenous promoter or under the strong GPD promoter, was also distributed in these vacuolar protrusions (Fig. 4B and Fig. S4A). When overexpressed, Ncr1-yeGFP behaved similarly to the endogenously expressed protein, forming NPC puncta that dissolved upon reduction of the AA concentration in the culture medium (Fig. S4B). These results indicate that NPC puncta correspond to vacuolar protrusions that are continuous with the main vacuolar membrane.

**Figure 4.**
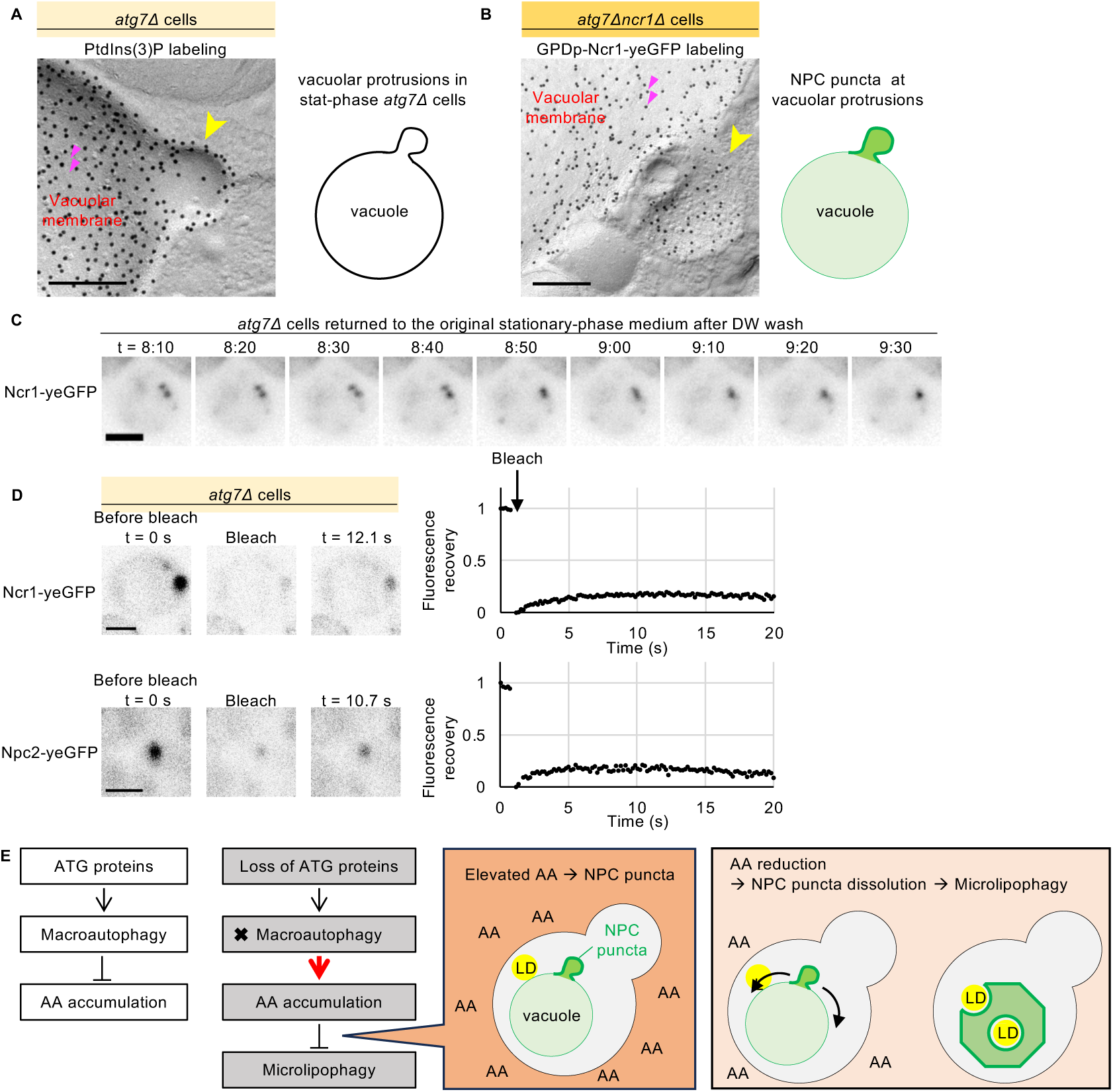
Liquid-like properties of NPC puncta formed in vacuolar protrusions. A) QF-FRL electron micrograph of stationary-phase *atg7*Δ cells expressing Ncr1-yeGFP showing a vacuolar protrusion (yellow arrowhead). The black dots represent gold colloid-labeled PtdIns(3)P (red arrowheads). Bar, 200 nm. B) QF-FRL electron micrograph of stationary-phase *atg7*Δ*ncr1*Δ cells overexpressing Ncr1-yeGFP. The yellow arrowhead shows a vacuolar protrusion, and the black dots represent gold colloid-labeled Ncr1-yeGFP (red arrowheads). Bar, 200 nm. C) Time-lapse images of stationary-phase *atg7*Δ cells expressing Ncr1-yeGFP washed with DW and subsequently cultured in their original medium. Bar, 2.5 μm. D) FRAP analysis of Ncr1-yeGFP and Npc2-yeGFP in *atg7*Δ cells. Time-lapse images show pre- and post-bleaching. Bar, 5 μm. The vertical axis indicates the relative fluorescence intensity (mean of >5 cells from a representative experiment; *n* = 2 independent experiments). E) Schematic: In stationary-phase *atg*Δ cells, extracellular AA accumulation induces NPC puncta formation. Upon AA reduction, the NPC protein distribution is immediately normalized, leading to the formation of vacuolar microdomains and the induction of microlipophagy.

Given this membrane continuity, the dissolution of NPC puncta upon AA reduction likely occurs through lateral diffusion rather than through vesicle-mediated transport or other energy-dependent processes. To test this idea, we examined whether NPC puncta dissolve even when active processes are inhibited. We found that NPC puncta dissolution in *atg7*Δ cells was not affected by ATP depletion (Fig. S4C), inactivation of the t-SNARE Vam3 in temperature-sensitive cells (Fig. S4D), or inhibition of actin and microtubule polymerization (Fig. S4E). These results are consistent with a model in which NPC proteins disperse from puncta largely through passive diffusion within a continuous vacuolar membrane system.

The rapid and reversible formation of NPC puncta suggests that they have liquid-like properties. Consistently, treatment with 1,6-hexanediol (1,6-HD) reversibly eliminated puncta formed by either Ncr1-yeGFP or Npc2-yeGFP (Fig. S4F), and Ncr1-yeGFP puncta underwent fusion (Fig. 4C). Fluorescence recovery after photobleaching (FRAP) analysis showed that both Ncr1-yeGFP and Npc2-yeGFP in puncta recovered to about 20% of their initial fluorescence (Fig. 4D), indicating that a fraction of NPC molecules remains exchangeable on this timescale. Consistent with this idea, bleaching a single punctum reduced the fluorescence of unbleached puncta in the same cell (Fig. S4G), suggesting molecular exchange between puncta. Together, these results indicate that NPC puncta formed in vacuolar protrusions are reversible clusters with liquid-like properties.

Furthermore, these clusters exhibit compositional selectivity. Unlike Ncr1-yeGFP and Npc2-yeGFP, Vph1-yeGFP, GFP-Pho8, and mRuby2-Cps1 did not show a punctate distribution in *atg7*Δ cells (Fig. S4H). Moreover, Npc2-yeGFP and Ncr1-yeGFP still formed puncta in *ncr1*Δ and *npc2*Δ cells, respectively, indicating that puncta formation does not require the other NPC protein (Fig. S4I). Together, these findings indicate that NPC proteins form selective, reversible clusters with liquid-like properties within distinct vacuolar protrusions.

## Discussion

Mechanistic understanding of microautophagy has lagged behind that of macroautophagy. A central question is whether the core ATG proteins, which are indispensable for macroautophagy, are also required for microautophagy.^3,4,20,21^ Here, we provide evidence that the apparent requirement for ATG proteins is indirect. In our working model (Fig. 4E), loss of macroautophagy in *atg*Δ cells leads to abnormal extracellular AA accumulation, reversible NPC puncta formation in vacuolar protrusions, disruption of vacuolar microdomains, and impaired microlipophagy. Importantly, reducing extracellular AA restores vacuolar microdomains and permits microlipophagy even in the absence of ATG proteins. These findings show that the stationary-phase microlipophagy defects in *atg*Δ cells is not a direct consequence of ATG loss, but instead results from metabolic dysregulation caused by macroautophagy deficiency.

This framework helps explain conflicting prior reports on the ATG dependence of microautophagy. Because AA, a byproduct of glucose fermentation, accumulates in the medium primarily after the diauxic shift,^32^ differences in growth phase and metabolic state would be expected to influence extracellular AA levels. This interpretation is consistent with reports showing that microlipophagy can occur in *atg*Δ cells during the post-diauxic shift phase,^23^ and that vacuolar microdomains are gradually lost thereafter.^15^ It may also explain why microlipophagy occurs during the ethanol-depleted phase of *atg*Δ cells, because extracellular AA would not be expected to accumulate during this phase due to the low level of glucose.^24,32^ Together, these observations suggest that apparent discrepancies among previous studies may reflect differences in extracellular AA levels.

The cause of abnormal AA accumulation in stationary-phase *atg*Δ cultures remains to be determined.^33^ One possible contributor is dysregulation of the acetaldehyde dehydrogenase Ald6, a known selective cargo of macroautophagy.^37,38^ In *atg*Δ cells, Ald6 might accumulate in the cytosol and promote excessive acetate production. Altered mitochondrial acetate metabolism may also contribute. For example, loss of Ach1, a mitochondrial succinyl-CoA:acetate CoA-transferase that converts acetate to acetyl-CoA, is associated with marked extracellular AA accumulation during the stationary phase.^39^ Although these possibilities remain speculative, they suggest that macroautophagy may help limit extracellular AA accumulation during stationary phase and thereby indirectly facilitate microlipophagy.

The rapid reformation of vacuolar microdomains and rescue of microlipophagy after AA reduction may reflect the physical properties of the AA-induced NPC puncta. Rather than forming static aggregates, these puncta behave as reversible clusters with liquid-like properties. Their fusion behavior, sensitivity to 1,6-HD, and rapid dissolution after AA reduction all support this interpretation. However, the molecular basis of puncta formation and the reason why NPC proteins in this punctate state do not support vacuolar microdomain formation remain unresolved. Npc2-yeGFP and Ncr1-yeGFP still formed puncta in *ncr1*Δ and *npc2*Δ cells, respectively (Fig. S4I), indicating that NPC puncta formation is not driven by direct binding between the two proteins. One possibility is that AA promotes the formation of a specialized physicochemical environment within vacuolar protrusions that favors partitioning of both proteins independently of direct interaction. Whether this reflects local changes in membrane organization, luminal composition, or both remains to be determined.

Although our study showed that exogenous AA reversibly suppresses microlipophagy in WT cells, it remains unclear whether endogenous fluctuations in AA regulate microlipophagy in a similar manner. Extracellular AA has been reported to exceed 50 mM in WT cells under certain conditions.^40^ This raises the possibility that AA-dependent disruption of vacuolar microdomains and impairment of microlipophagy may also occur in WT cells under physiological conditions, thereby contributing to the previously reported shortening of chronological lifespan by AA.^32^

In summary, our study addresses a long-standing debate by providing evidence that ATG proteins are not required for the execution of stationary-phase microlipophagy. We therefore propose that macroautophagy supports stationary-phase microlipophagy indirectly by limiting AA accumulation and thereby preventing AA-induced NPC clustering and loss of vacuolar microdomains.

### Limitations of this study

A key limitation of this study is that the precise mechanism by which AA alters NPC protein behavior and impairs their role in vacuolar microdomain formation has not yet been defined. Our data suggest that Ncr1 and Npc2 can form clusters independently of each other. However, whether this clustering is driven by additional proteins, altered lipid composition, or local pH changes remains to be determined.

## Materials and methods

### Yeast

The *S. cerevisiae* strains used in this study are derivatives of SEY6210 and are listed in Table S1. Gene deletions were performed using PCR-based homologous recombination with pFA6a plasmids as templates following standard protocols.^41^

Cells were cultured in 100 mL flasks containing 10 mL of SD + CA + ATU medium (pH 6.0), composed of 0.17% yeast nitrogen base without amino acids and ammonium sulfate (BD Difco), 0.5% ammonium sulfate (Wako Pure Chemical Industries), 0.5% casamino acids (BD Difco), 0.004% adenine (Wako), 0.004% tryptophan (Wako), 0.002% uracil (Wako), and 2% dextrose (Wako), in a shaking incubator at 30°C. For log-phase cultures, cells were grown starting from an optical density at 600 nm (OD_600_) of 0.15 until they reached an OD_600_ of 0.6–0.8. Stationary-phase cultures were obtained by growing the cells for 2–4 days. Culture media from WT and *atg*Δ cells in the stationary phase were collected by centrifugation at 2,000 × *g* for 1 min, filtered through 0.22 μm filters, stored at −80°C, and used for subsequent experiments. The *vam3* temperature-sensitive mutant was cultured at the non-permissive temperature of 37°C for 3 h prior to analysis.

### Chemical treatment

Cells were treated with culture medium supplemented with 10% 1,6-HD (Wako) for 10 min at RT. To deplete ATP, cells were treated with 2% 2-deoxyglucose and 3 mM sodium azide for 10 min in their medium. To assess the influence of the cytoskeleton on NPC puncta dissolution, cells were incubated with their medium supplemented with either 1 μM cytochalasin D (Sigma‒Aldrich) for 30 min or 10 μg/mL nocodazole (Sigma‒Aldrich) for 4 h.^28^ After treatment, cells were washed with DW containing the respective inhibitors at the same concentration.

### AA treatment

The concentration of AA in the culture supernatants was quantified using an acetate colorimetric assay kit (MAK086, Sigma‒Aldrich) according to the manufacturer’s instructions. AA was added to WT cultures to reach a final concentration of 44 mM. For experiments requiring AA reduction, cells were harvested by centrifugation at 2,000 × *g* for 1 min and resuspended in fresh medium, DW, or culture medium from stationary-phase WT cells. The pH of the medium was adjusted by adding NaOH. NaCl was added at an equivalent molar concentration as a control for ionic strength. Other organic acids (fumaric, succinic, citric, and malic acids) were tested at molar concentrations equal to or greater than that of AA. Acetate buffer (50 mM, pH 3.6) was used as a wash solution to replace medium components while maintaining AA.

### Fluorescence microscopy

Yeast cells expressing yeGFP- or mRFP-tagged proteins were mounted on glass slides and covered with coverslips. Images were acquired using an Axiovert microscope equipped with a 100× Apochromat objective lens (Carl Zeiss) and the acquisition software ZEN (Carl Zeiss). Images were processed using Fiji/ImageJ and Adobe Photoshop.

To label LDs, yeast cells were incubated with 1 μg/mL BODIPY 493/503 (Thermo Fisher Scientific) for 10 min before observation. The vacuolar membrane was stained with 400 μM FM4-64 (Invitrogen) for 20 min at 30°C on the day before observation. Cells were resuspended in their original medium after being washed twice with DW.

FRAP experiments were conducted with a laser scanning confocal microscope [LSM780 (Zeiss) or FV3000RS (EVIDENT)]. NPC puncta were photobleached at maximum laser power. Fluorescence intensities were quantified using Fiji/ImageJ and normalized to the prebleaching intensity (set to 1.0) and the minimum postbleach intensity (set to 0).

### Quick-freezing and freeze-fracture replica preparation

Ultrastructural analysis using QF-FRL to visualize NPC puncta in atg7Δ cells was chosen because aldehyde fixatives were ineffective for NPC puncta, as reported for certain liquid droplets.^42^ Yeast cells were quickly frozen via high-pressure freezing using an HPM 010 or HPM 100 high-pressure freezing machine (Leica Microsystems). Cells were harvested by centrifugation at 2,000 × *g* for 1 min, sandwiched between a 20 μm-thick copper foil and a flat aluminum disc (Engineering Office M. Wohlwend), and frozen according to the manufacturer’s instructions.^28^

For freeze-fracture, the frozen samples were transferred to the cold stage of an ACE900 instrument (Leica Microsystems) and fractured at −102°C under a vacuum of ∼1 × 10^−6^ mbar. Replicas were made via electron beam evaporation in three steps: (1) carbon (6 nm thick) at a 90° angle to the sample surface; (2) platinum-carbon (2 nm thick) at a 45° angle; and (3) carbon (10 nm thick) at a 90° angle.

Thawed replicas were treated with 2.5% SDS in 0.1 M Tris-HCl (pH 8.0) at 60°C overnight. To remove the cell wall, replicas were treated for 20 min or 2 h (for protein labeling or lipid labeling, respectively) at 30°C with 0.1% Westase (Takara Bio) in McIlvaine buffer (pH 6.0) supplemented with 10 mM EDTA, 30% fetal bovine serum, and a protease inhibitor cocktail. After enzyme treatment, the replicas were further treated with 2.5% SDS and stored in 50% glycerol in phosphate-buffered saline (PBS) at −20°C until use.

### Replica labeling

PtdIns(3)P was labeled as described previously.^43^ Briefly, SDS-treated freeze-fracture replicas were incubated at 4°C overnight with GST-tagged p40phox PX domain (10 ng/mL) in PBS containing 1% bovine serum albumin (BSA), followed by incubation with rabbit anti-GST antibody (5 μg/mL, Bethyl Laboratories) and then with colloidal gold (10 nm)-conjugated protein A (PAG10; University of Utrecht), each for 40 min at 37°C in PBS supplemented with 1% BSA. For Ncr1-yeGFP labeling, replicas were incubated with a rabbit anti-GFP antibody (provided by Dr. Masahiko Watanabe of Hokkaido University) followed by PAG10. The labeled replicas were placed on Formvar-coated EM grids and observed with a JEM-1011 (JEOL) operated at 100 kV. Digital images were captured using a CCD camera (Gatan).

### Western blotting

Yeast cell extracts were prepared using lithium acetate and sodium hydroxide.^44^ The precipitated samples were dissolved in SDS sample buffer (2% SDS, 10% glycerol, and 60 mM Tris-HCl, pH 6.8) and centrifuged. The supernatant was subjected to SDS‒PAGE, electrotransferred to a PVDF membrane, and analyzed by Western blotting using SuperSignal West Dura Extended Duration Substrate (Thermo Fisher Scientific). Images were captured with a Fusion Solo S instrument (Vilber Lourmat) and analyzed using Fiji/ImageJ.

### Cytosolic pH measurement

The cytosolic pH was measured using the ratiometric fluorescent probe pHluorin or the Rosella biosensor (a fusion of pHluorin and DsRed.T3).^45^ Fluorescence intensity ratios were calculated for individual cells. Calibration curves were generated by equilibrating cells in citric acid-Na_2_HPO_4_ buffers of known pH containing 2 mM 2,4-dinitrophenol (DNP), 2% 2-deoxyglucose, and 3 mM sodium azide.

### Statistical analysis

Data are presented as mean ± SEM from at least three independent experiments unless stated otherwise. Student’s t-test or Mann–Whitney U test was used for comparisons between two groups, as indicated in the figure legends. Statistical significance was set at *p* < 0.05 (*) and *p* < 0.01 (**) where indicated. The number of cells analyzed per condition is specified in the figure legends (typically more than 50 cells per experiment).

### Box plots

Box plots were prepared using BoxPlotR (http://boxplot.tyerslab.com/). The centerlines show the medians, the box boundaries indicate the 25th and 75th percentiles, the whiskers delineate the maximum and minimum data points that are no more than 1.5 times the interquartile range, and the dots represent individual data points. The average is shown by +.

## Acknowledgments

We thank Dr. Masahiko Watanabe (Hokkaido University) for the anti-GFP antibody; Dr. Yui Jin (The Nippon Dental University) for GFP-Pho8, mRuby2-Cps1, and Kog1-3×GFP; Dr. Rodney Devenish and Dr. Carlos Rosado (Monash University) for the pH biosensor Rosella; Ms. Asami Maeda and Ms. Hiroko Osakada-Iwamoto (Juntendo University) for excellent technical assistance. We also thank Dr. Yoshinori Ohsumi (Tokyo Institute of Technology), Dr. Yasuyoshi Sakai and Dr. Masahide Oku (Kyoto University) for helpful suggestions. This study was supported by JSPS KAKENHI (grant numbers 22K06818 and 22H04654 to T.T. and grant number 22H00446 to T.F.) and JST CREST (grant number JPMJCR20E3 to N.N.) and by a grant from the Research Institute for Diseases of Old Age, Juntendo University Graduate School of Medicine (to T.T.).

## Author contributions

T.T., M.F., and N.N. conducted the experiments; T.T. and T.F. designed the experiments; and T.T., N.N., and T.F. wrote the manuscript.

## Declaration of generative AI and AI-assisted technologies in the writing process

During the preparation of this work, the authors used ChatGPT (OpenAI) for English grammar and language editing only. The authors reviewed and revised all output and take full responsibility for the final content of the manuscript.

---

**Figure S1.**
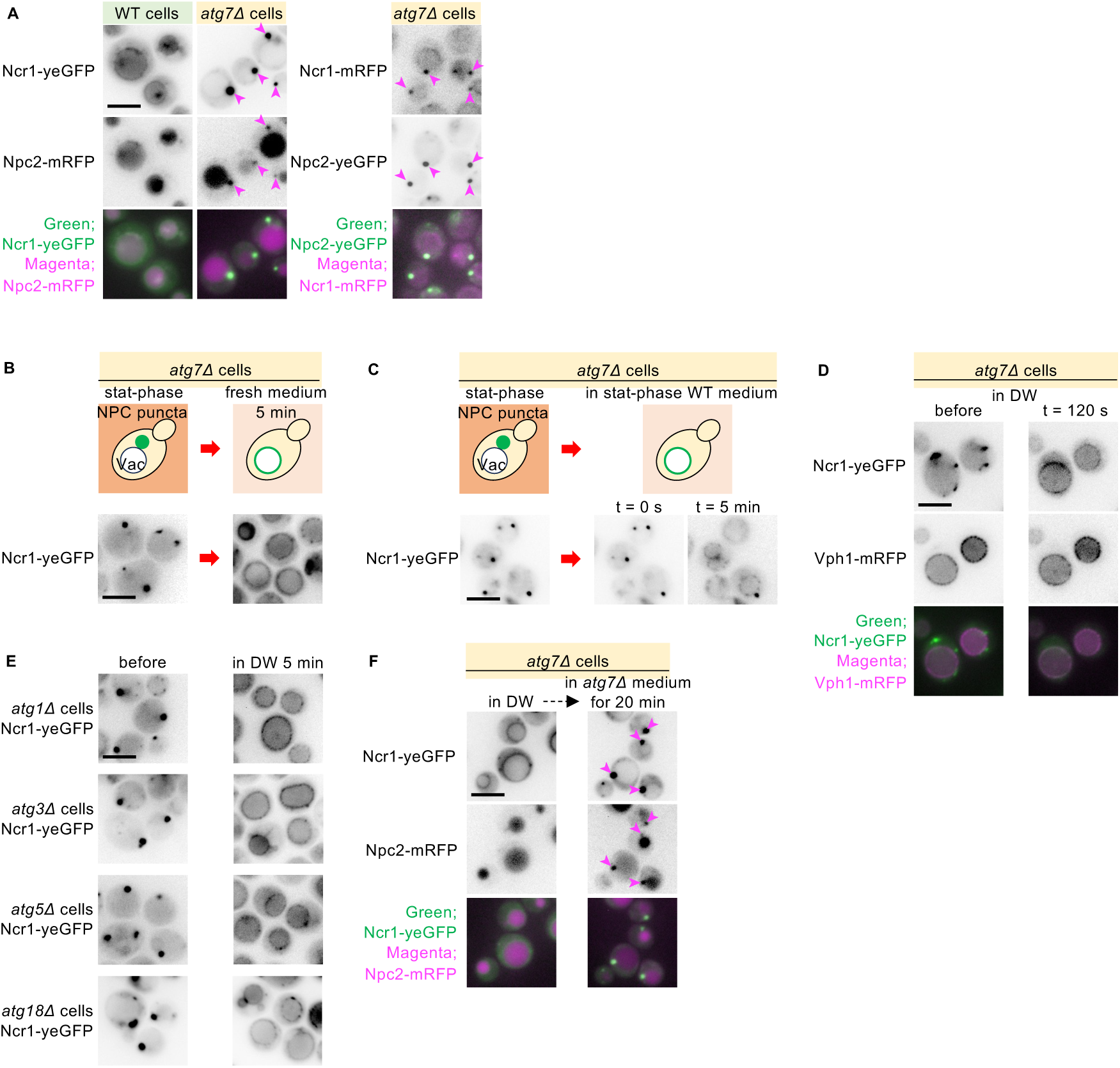
NPC puncta formation and dissolution. A) Colocalization of Ncr1 and Npc2 at NPC puncta (arrowheads) in stationary-phase *atg7*Δ cells. Bar, 5 μm. B) Dissolution of NPC puncta in stationary-phase *atg7*Δ cells after 5 min culture in fresh medium. Bar, 5 μm. C) Time-lapse images of Ncr1-yeGFP in stationary-phase *atg7*Δ cells that were cultured in the stationary-phase WT medium. Bar, 5 μm. D) Time-lapse images of Ncr1-yeGFP and Vph1-mRFP in stationary-phase *atg7*Δ cells upon DW wash. Bar, 5 μm. E) NPC puncta dissolution in stationary-phase *atg1*Δ, *atg3*Δ, *atg5*Δ, and *atg18*Δ cells after washing with DW. Bar, 5 μm. F) Ncr1-yeGFP and Npc2-mRFP in stationary-phase *atg7*Δ cells that were washed with DW and returned to their own stationary-phase medium. The reformed NPC puncta harboring both Ncr1-yeGFP and Npc2-mRFP are shown by arrowheads. Bar, 5 μm.

**Figure S2.**
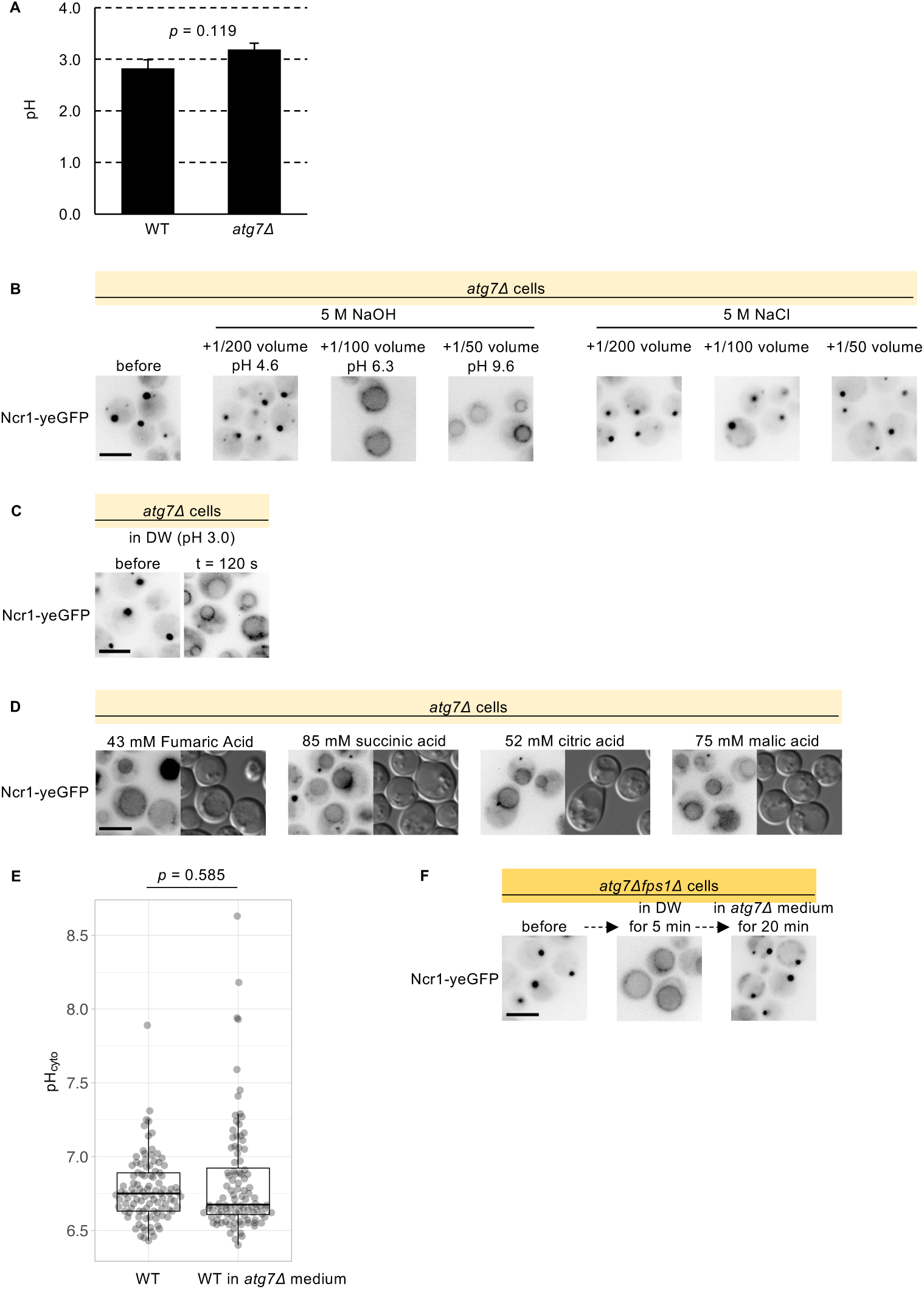
NPC protein distribution under several conditions. A) pH values of the medium of WT and *atg7*Δ cells in the stationary phase. The mean ± SEM of three independent experiments is shown. Student’s t-test. B) Representative images of stationary-phase *atg7*Δ cells expressing Ncr1-yeGFP cultured in the medium supplemented with NaOH or NaCl for 5 min. Bar, 5 μm. C) Time-lapse images of stationary-phase *atg7*Δ cells expressing Ncr1-yeGFP after transfer to HCl-acidified DW (pH 3.0). NPC puncta dissolved within 2 min. Bar, 5 μm. D) Representative images of stationary-phase *atg7*Δ cells expressing Ncr1-yeGFP after washing with organic acids, showing NPC puncta dissolution. Bar, 5 μm. E) Cytosolic pH values of stationary-phase WT expressing a fluorescent pH biosensor (Rosella) before and after culturing in *atg7*Δ medium for 1 h. Data from 100 cells per condition (50 cells in each of two independent experiments) were pooled and are shown as boxplots. Statistical significance was evaluated using the Mann–Whitney U test. F) Representative images of stationary-phase *atg7*Δ*fps1*Δ cells expressing Ncr1-yeGFP washed with DW and then cultured in the *atg7*Δ medium. Bar, 5 μm.

**Figure S3.**
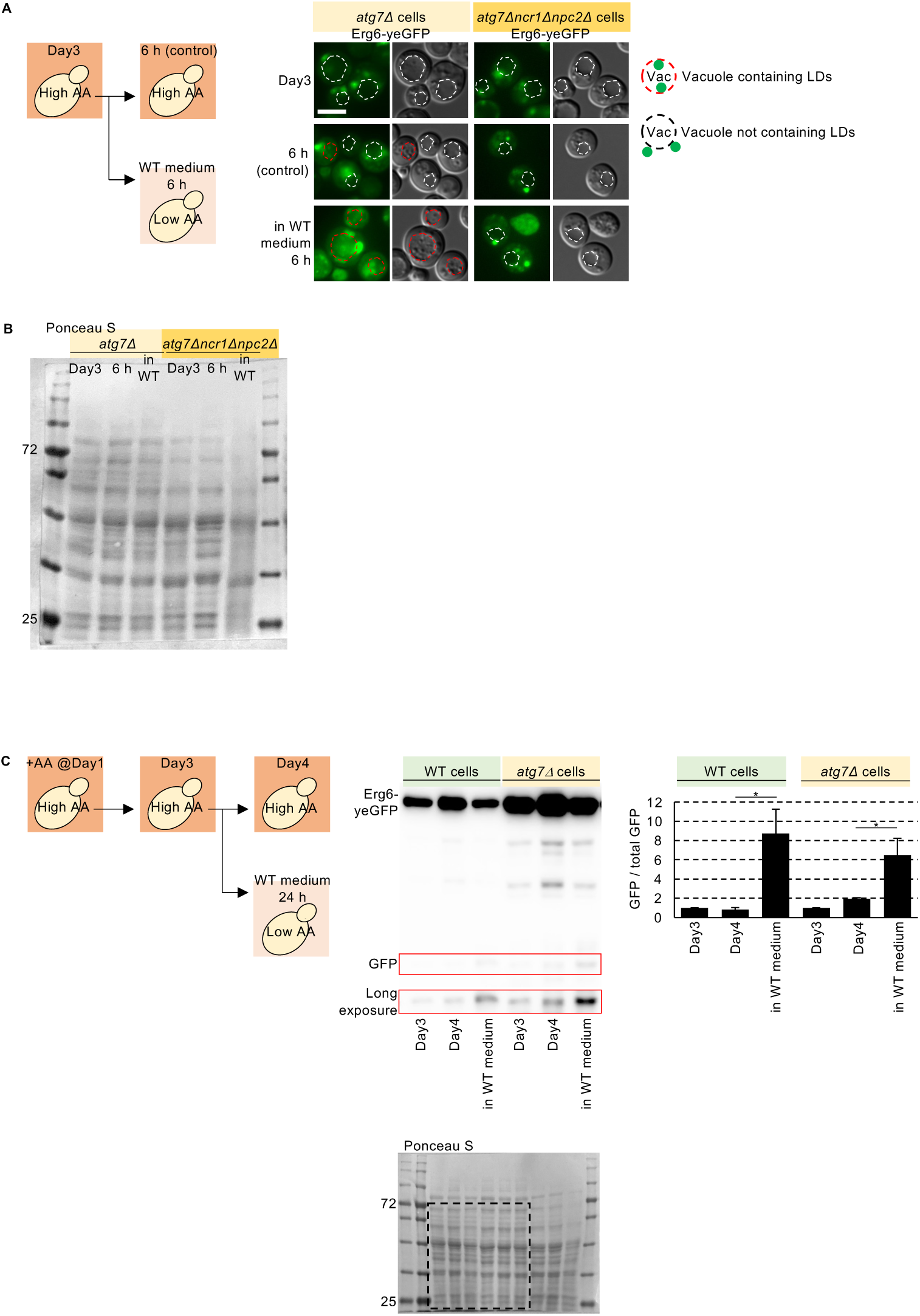
Vacuolar microdomain formation and stationary-phase microlipophagy in *atg*Δ cells. A) Representative images of *atg7*Δ and *atg7*Δ*ncr1*Δ*npc2*Δ cells expressing Erg6-yeGFP before and after culture in the stationary-phase WT medium for 6 h. The red dotted lines indicate vacuoles incorporating LDs, whereas the white dotted lines indicate vacuoles without LDs. Bar, 5 μm. B) Ponceau S staining corresponding to Fig. 3D. C) Vacuolar degradation of the LD marker Erg6-yeGFP in WT cells and *atg7*Δ cells was assessed on Days 3 and 4. On Day 0, cells were diluted in fresh medium at OD_600_ = 0.15, and AA was added to the medium on Day 1. On Day 3, half of the cells were transferred to the stationary-phase WT medium without AA supplementation for an additional 24 h. The relative ratio of free GFP to full-length protein was measured. The mean ± SEM of three independent experiments is shown. Student’s t-test (Day 4 vs. Day 4 in WT medium). *; *p* < 0.05.

**Figure S4.**
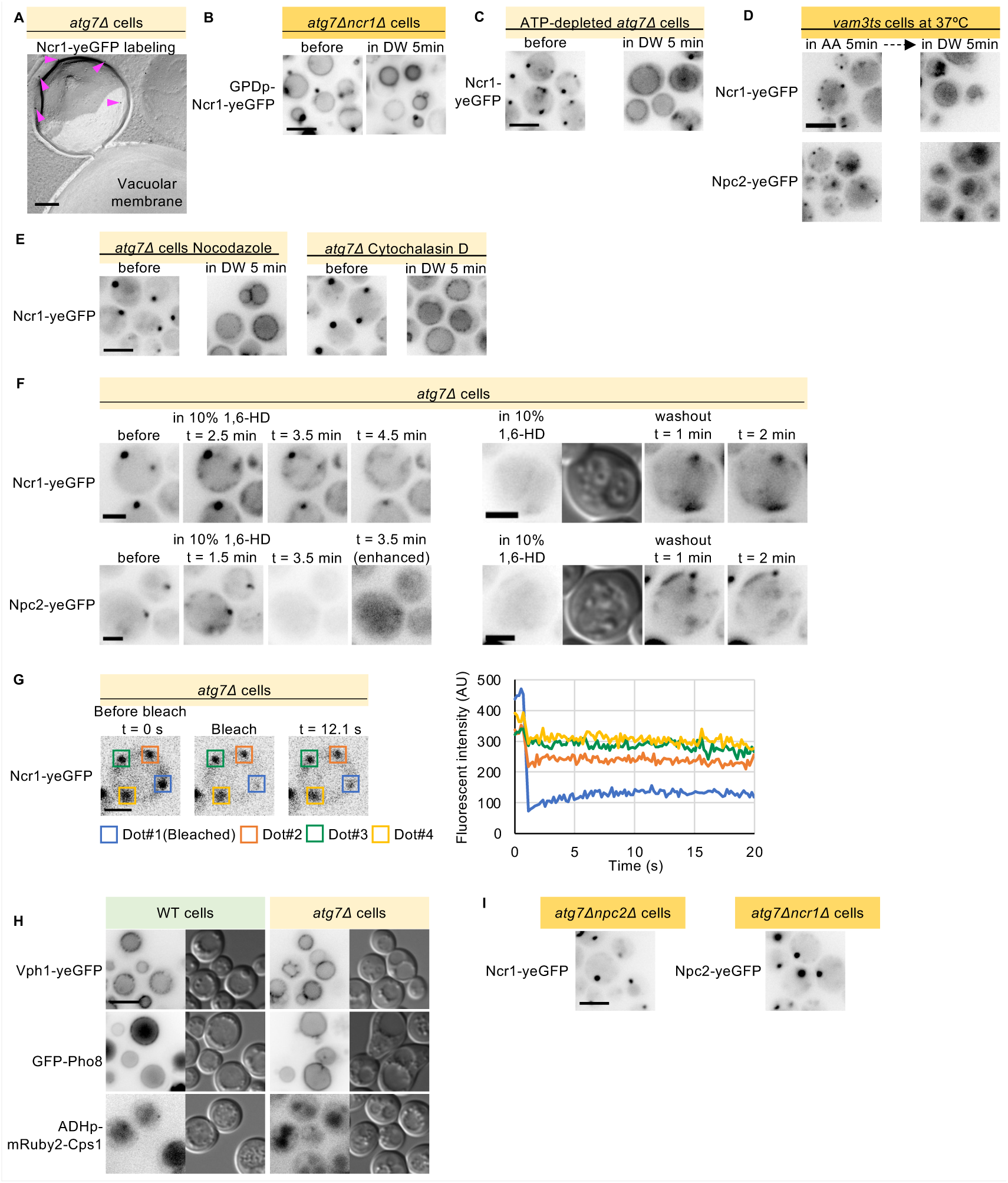
NPC puncta formed in vacuolar protrusions. A) QF-FRL electron micrographs of stationary-phase *atg7*Δ cells expressing Ncr1-yeGFP. The black dots represent gold colloid-labeled Ncr1-yeGFP (arrowheads). Bar, 200 nm. B) Representative images of stationary-phase *atg7*Δ*ncr1*Δ cells overexpressing Ncr1-yeGFP. Bar, 5 μm. C) Representative images of stationary-phase *atg7*Δ cells expressing Ncr1-yeGFP treated with 2% 2-deoxyglucose and 3 mM NaN_3_ for 10 min, and then washed with DW containing 2% 2-deoxyglucose and 3 mM NaN_3_ for 5 min. Bar, 5 μm. D) Representative images of stationary-phase *vam3*-ts cells expressing Ncr1-yeGFP cultured at the non-permissive temperature of 37°C for 3 h and maintained at 37°C during exposure to and washout of AA, showing NPC puncta formation and dissolution. Bar, 5 μm. E) Representative images of stationary-phase *atg7*Δ cells expressing Ncr1-yeGFP treated with 10 μg/mL nocodazole for 4 h or 1 μM cytochalasin D for 30 min, followed by washing with DW containing 10 μg/mL nocodazole or 1 μM cytochalasin D for 5 min. Bar, 5 μm. F) Time-lapse images of *atg7*Δ cells showing the dissolution and reformation of NPC puncta during treatment with 1,6-HD and subsequent washout of 1,6-HD, respectively. Bar, 2.5 μm. G) FRAP analysis of Ncr1-yeGFP in *atg7*Δ cells with multiple NPC puncta. Photobleaching of a single punctum (dot #1) results in decreased fluorescence in the other puncta. Bar, 2.5 μm. The vertical axis of the graph indicates the fluorescence intensity of Ncr1-yeGFP. H) Micrographs of stationary-phase WT and *atg7*Δ cells expressing Vph1-yeGFP, GFP-Pho8, mRuby2-Cps1. Bar, 5 μm. I) Representative images of stationary-phase *atg7*Δ*npc2*Δ cells expressing Ncr1-yeGFP and *atg7*Δ*ncr1*Δ cells expressing Npc2-yeGFP showing NPC puncta formation. Bar, 5 μm.

**Movie 1.**

Six-panel DIC time-lapse movie of *atg7*Δ and WT cells under the indicated AA conditions. The top row shows *atg7*Δ cells and the bottom row shows WT cells. From left to right, cells are shown on Day 3 after AA supplementation on Day 1, on Day 4 after continuous AA supplementation from Day 1, and on Day 4 after AA reduction on Day 3. Vacuoles are largely devoid of internal vesicles before AA reduction, whereas numerous vesicles undergoing Brownian motion are visible within vacuoles after AA reduction. Images were acquired at 0.5 s intervals.

**Movie 2.**

Fluorescence time-lapse movie of *atg7*Δ cells on Day 4 after AA reduction on Day 3. LDs stained with BODIPY 493/503 are visible within the vacuoles. Images were acquired at 0.5 s intervals.

**Movie 3.**

Fluorescence time-lapse movie of WT cells on Day 4 after AA reduction on Day 3. LDs stained with BODIPY 493/503 are visible within the vacuoles. Images were acquired at 0.5 s intervals.

**Table S1.**
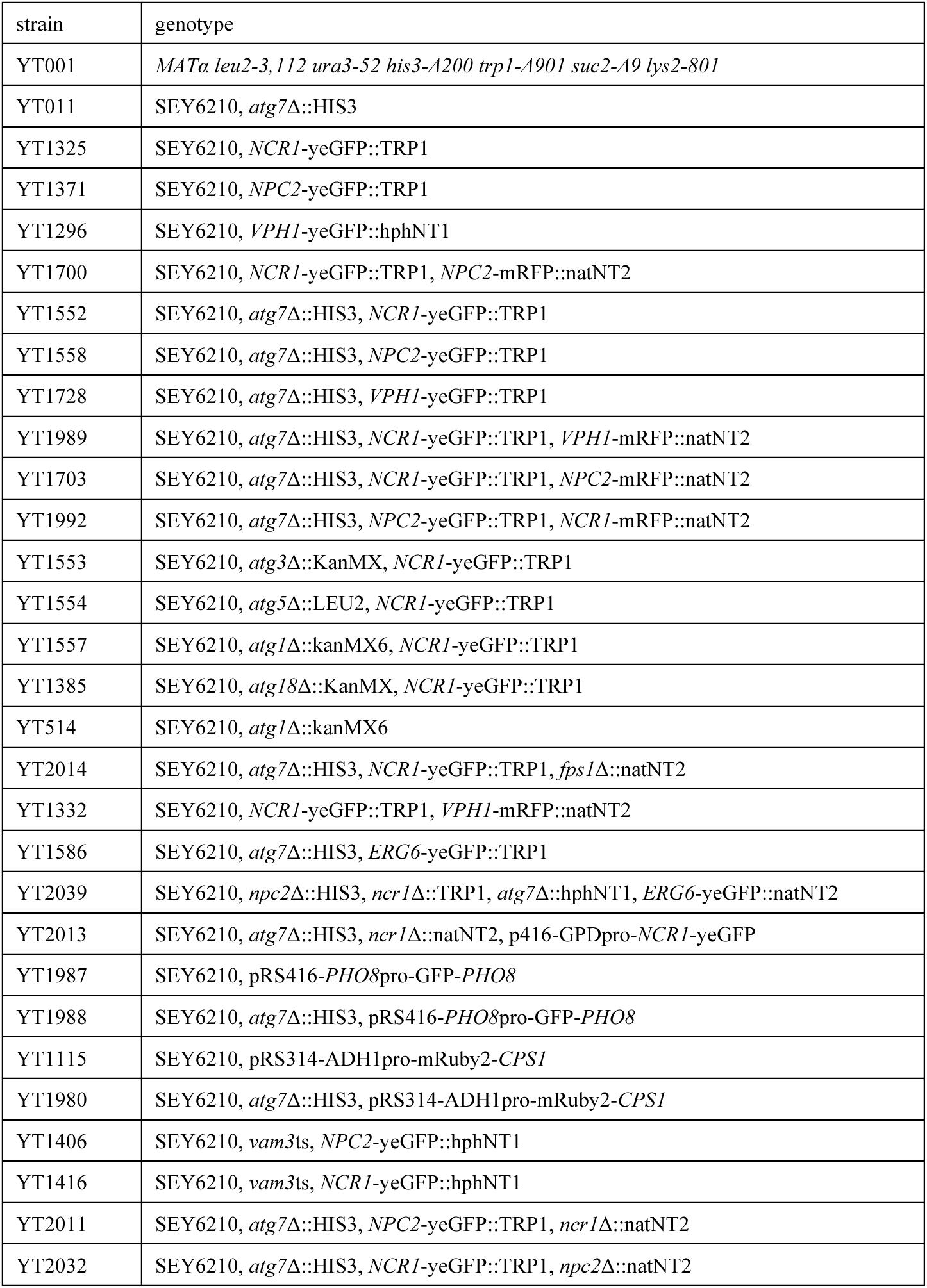
Yeast strains used in this study.

